# Microdiversity in marine pelagic ammonia-oxidizing archaeal populations

**DOI:** 10.1101/2024.04.23.590705

**Authors:** Pablo Suárez-Moo, Jose M. Haro-Moreno, Francisco Rodriguez-Valera

**Affiliations:** Evolutionary Genomics Group, División de Microbiología, Universidad Miguel Hernández, Apartado 18, San Juan 03550, Alicante, Spain

**Keywords:** Ammonia-oxidizing archaea, metagenomic recruitment, long-read metagenomics, flexible genomic islands, *Nitrosopelagicus brevis*, *Nitrosopumilus catalinensis*

## Abstract

The knowledge of the different population-level processes operating within a species, and the genetic variability of the individual prokaryotic genomes, is key to understanding the adaptability of microbial populations. Here, we characterized the flexible genome of ammonia-oxidizing archaeal (AOA) populations using a metagenomic recruitment approach and long-read (PacBio HiFi) metagenomic sequencing. In the lower photic zone of the western Mediterranean Sea, the genomes *Nitrosopelagicus brevis* CN25 and *Nitrosopumilus catalinensis* SPOT1 had the highest recruitment values among AOA. These two complete genomes were used to analyze the diversity of flexible genes (variable from strain to strain) by analyzing the long-reads located within the flexible genomic islands (fGIs) identified through their under-recruitment. Both AOA genomes had a large fGI involved in the glycosylation of exposed structures, highly variable and rich in glycosyltransferases. *N. brevis* had two fGIs related to the transport of phosphorus and ammonium respectively. *N. catalinensis* had fGIs involved in phosphorus transportation and metal uptake. A flexible genomic island (fGI5) previously reported as “unassigned function” in *N. brevis* could be associated with defense. These findings demonstrate that the microdiversity of marine microbe populations, including AOA, can be effectively characterized using an approach that incorporates new third-generation sequencing metagenomics.

## INTRODUCTION

One remarkable discovery of prokaryotic population genomics is the large diversity, particularly in the gene pools of individuals (or strains) belonging to the same species and coming from the same sample (Van Rossum et al. 2020). This is particularly remarkable when the sample is relatively small and belongs to a rather homogeneous habitat, such as the water column of the oligotrophic off-shore ocean (Haro-Moreno et al. 2018; Haro- Moreno et al. 2024). This diversity was hidden by the low culturability of marine microbes and had to wait until metagenomic approaches were developed (Cuadros-Orellana et al., 2007; Coleman et al., 2006).

There are two possible non-exclusive explanations for this specific example of the “plankton paradox”. One is the diversity of substrates in the case of heterotrophic (frequently, photoheterotrophic) that bacteria can find in the ocean. It has been calculated that ca. 100.000 different chemical formulas of organic compounds (not counting isomers) are found as DOM in the water column of the open ocean (Riedel and Dittmar 2014). This vast figure explains that large gene pools of transporters and collateral metabolic pathways would be required to degrade such an enormous diversity of compounds and close the carbon cycle. Another (complementary) explanation suggested by some of us is that high intrapopulation diversity is maintained by the Constant Diversity (CD) equilibrium reached with the cognate phage populations (Rodriguez-Valera et al. 2009). Both processes could be concurrent and also need each other to keep the ecosystem functioning efficiently.

Ammonia-oxidizing archaea (AOA) or Thaumarchaeota represent an important group associated with the nitrogen cycle (ammonia oxidation and primary productivity) in oligotrophic marine environments (Zheng et al. 2024). Recent population genomic studies in AOA have reported the functions and metabolic potential associated with adaptation to different specific environments such as marine sediments (Kerou et al. 2021), freshwater lakes (Klotz et al. 2022; Ngugi et al. 2023), and marine water column (Haro-Moreno et al. 2018). The analysis of the flexible genome through metagenomic analysis has been classically hampered by the poor assembly of short reads covering these regions. Hence we have used a long read (PacBio CCS) run to provide a better understanding of the AOA microdiversity.

Here we have analyzed a single population of two AOA species (*Nitrosopelagicus brevis* and *Nitrosopumilus catalinensis*) that were predominant in a microbial community found 80 m deep in the water column (lower photic zone) of off-shore Western Mediterranean at a station that has been extensively studied before by both short and long-read metagenomics (Haro-Moreno et al. 2018; Haro-Moreno et al. 2021). Given that these are chemolithotrophic microbes that have a major single energy and biomass-generating substrate (ammonia or urea and CO_2_) it was expected that less intrapopulation diversity would be found and certainly much less than in the case of heterotrophic microbes.

## RESULTS

We sequenced a single sample retrieved from ∼75 m deep at off-shore Mediterranean waters while the water column was stratified (early October). Previous work indicated that AOAs were very abundant at this depth where ammonia is relatively plentiful (Haro- Moreno et al. 2018). One metagenomic sample was retrieved and sequenced by both Illumina (29 Gbp) and PacBio Sequel II CCS15 (9.2 Gbp). A total of 194.4 and 1.9 million high-quality and filtered reads were obtained (Table A1). The community structure revealed by either fragment (∼150 nucleotides) of the 16S rRNA gene (Illumina short reads, SR) or by PacBio long reads (LR) near complete 16S rRNA gene (∼1,400 nucleotides) was very similar (Figure A1), confirming the high abundance of AOA in the sample (ca. 9% of the short read 16S fragments and 12% of the PacBio reads containing 16S rRNA genes).

### A metagenomic strategy to determine flexible genomic islands

Our strategy to obtain data on the local pangenome of the abundant species of AOA was based on differential recruitment (Rodriguez-Valera et al., 2009). Briefly, using a reference genome that is assumed to have close relatives (members of the same species) in the sample, the regions that recruit significantly fewer reads are considered “metagenomic islands”, i.e. regions of the reference genome with little or no presence in the metagenome and likely belonging to the flexible genome of the species. Since no culture has been retrieved from this location (Mediterranean Sea), we identified all the AOA 16S rRNA genes obtained in the LR metagenome to check if some corresponded to the available AOA genomes. A total of 440 near-complete (>1,400 nucleotides) 16S rRNA gene sequences associated with Thaumarcheota were extracted from the LRs and clustered in 15 representative sequences (>97% sequence identity) (Figure A1B). The *Nitrosopelagicus* and *Nitrosopumilus* genera had the largest number of 16S rRNA sequences that clustered within them with 312 and 98, respectively. In addition, 11 sequences appeared related to the pSL12 clade later described as the “heterotrophic marine thaumarchaea” (Aylward and Santoro 2020). Sequences very similar (>97% identity) to genomes of *Nitrosopelagicus brevis* CN25 and *Nitrosopumilus catalinensis* SPOT01 cultures were abundantly represented with 187 (43%) and 98 (23%) sequences, respectively (Figure A1B). Both strains are Pacific Ocean isolates, the first from offshore waters and the latter from more coastal regions, but the high similarity indicates that both species are well represented in our Mediterranean deep-photic-zone.

Next, we recruited these genomes against the SR (Illumina) metagenome showing that indeed very similar microbes were present in the Mediterranean sample (Table A2 and Figure 1). Although both genomes recruited largely at 100% similarity, *N. brevis* CN25 seems to have closer relatives in its local population while *N. catalinensis* SPOT01 had a dominant lineage different from the Pacific SPOT01 genome (Figure 1). However, we decided to use both reference genomes. Mediterranean populations revealed by metagenomic recruitment five and four metagenomic islands for *N. brevis* CN25 and *N. catalinensis* SPOT01, respectively (Figure 1, Tables A3 and A4).

**Figure 1.**
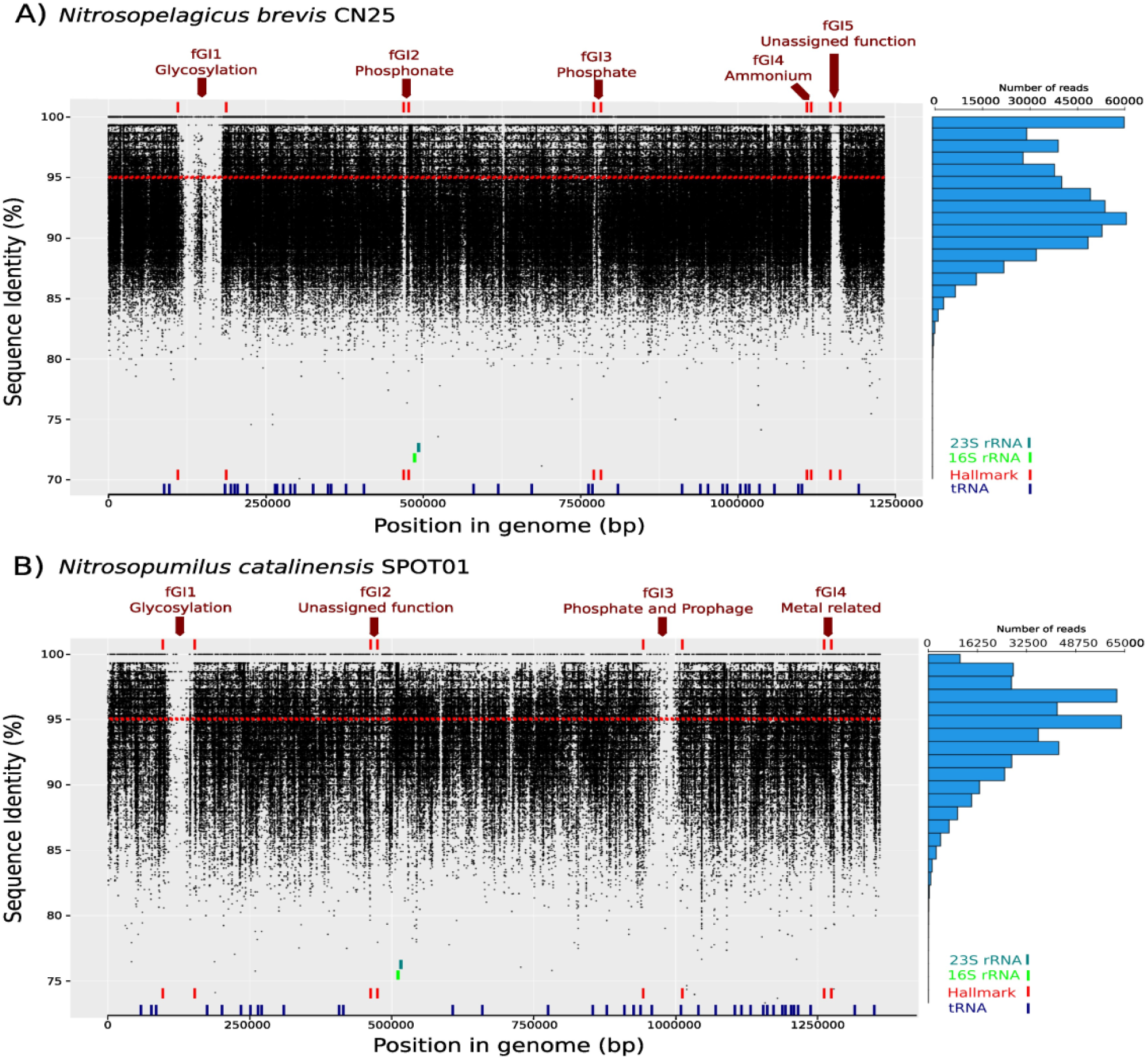
Flexible genomic islands (fGIs) and metagenomic recruitment of *Nitrosopelagicus brevis* CN25 (A) and *Nitrosopumilus catalinensis* SPOT01 genomes (B). The reference genomes were recruited against the Illumina (SR) dataset from the lower photic zone. The location of the fGI is shown at the top of each recruitment plot. The hallmarks (red lines up), 16S (green lines), and 23S (gray lines) rRNA gene, and tRNA (Blue lines) are shown at the lower part.

To reconstruct the regions found in different cells of the population (fGIs) we used the LRs from the PacBio metagenome. We selected hallmark genes (HG) that were located at or near the limit of the genomic island (Figure 1) and used them as bait to recover long reads that covered the hallmark plus the variable region. In addition, to expand the information about the different versions (gene content) of the fGIs we overlapped LRs that covered the region at a range of 97-100% similarity with each other (for details see Figures A2, A3-A6).

### Glycosylation-related fGIs

Both AOA species have a typical fGI related to glycosylation. The largest island in CN25 had been already identified by comparing the CN25 genome with several metagenomic datasets from marine environments, including the Global Ocean Sampling (GOS) (Santoro et al. 2015) (Figure 1A and Table A3) and our results appear to largely overlap those described there. The glycosylation fGI was the largest in both genomes (89,958 bp and 51,740 bp for CN25 and SPOT01, respectively). Glycosylation gene clusters of putatively exposed structures such as the lipopolysaccharide of Gram-negative bacteria are the most typical kind of fGI found in prokaryotes (Rodriguez-Valera, et al. 2016). They were found in halophilic archaea (Cuadros-Orellana et al., 2007) where they were proven to contain the major cell-surface glycoprotein (Martin-Cuadrado, et al. 2015). In bacteria, there are several depending on the phyla, the O-chain of the Gram-negative lipopolysaccharide being the most widely studied. In the case of CN25, the specific function of the glycosylation genes is unknown, as is the major component of the cell envelope. However, given the large numbers of glycosyltransferases and other genes related to polysaccharide synthesis and export, we can speculate that the genes are associated with the glycosylation of cell envelope components, and/or the synthesis of extracellular polysaccharides. In the case of *Nitrosopumilus maritimus* SCM1, the first genome of marine AOA known, the cell envelope is made up of a highly organized proteinaceous surface layer (S-layer) with p6 symmetry (Qin et al. 2017). Electron microscopy analysis showed a rather thick cell wall of ca. 20 nm and it is likely that the S-layer protein is glycosylated (Zhou et al. 2023). Both *N. brevis* and *N. catalinensis* genomes have ORFs that annotate as S-layer protein: ORF_1385 (1,313 bp) and ORF_1574 (1,316 bp) respectively, both located far from the fGI glycosylation island (Tables A3 and A4). On the other hand, no gene annotated as S-layer protein was found in any of the versions of this island (see below). For now, we can hypothesize that fGI1 codes for the glycosylation of an S-layer cell wall or an independent capsular polysaccharide. In *N. maritimus* SCM1, a similar glycosylation gene cluster was not found. However, six glycosyltransferase genes were found dispersed in the genome (ORF_63, ORF_118, ORF_168, ORF_252, ORF_464, ORF_624), the predicted S-layer proteinaceous component gene is located elsewhere (ORF_1822) (Table A5).

The specific gene cluster found at fGI1 in the strain CN25 could not be found in our metagenomes indicating that likely the clonal lineage represented by this strain is not present in our sample. However, we found several reads overlapping the hallmarks (HGs) (Figure 2A) and getting into the fGI. We have found a relatively low diversity of this gene cluster judging by the repeated finding of the same genes or highly similar. Thus, using the left HG one single version was found repeated 28 times and another 21 (Figure 2A). The right-hand end seemed to be more evenly distributed but still all versions were found more than once (ranging from 2 to 8 LRs by version). Contrastingly, for SPOT01 we did recover many genes similar to the island of the Pacific isolate genome (upper part of Figure 2B). Thus ca. 5 Kb of nearly identical reads to the left-hand side of the Pacific isolate was recovered 27 times and a smaller part (ca. 3 Kb) of the right-hand side was recovered three times. However, the most common version of this fGI (36 LRs) was not similar to the one of the Pacific Ocean isolate, with a deletion of most of the ORFs present in the SPOT01 genomic island (94%) (Figure 2B, lower part). Either for *N. brevis* or for *N. catalinensis,* the diversity of sequence of the glycosylation fGIs was much lower than that found in other microbes (Coleman et al. 2006; Haro-Moreno et al. 2024). However, in both cases, there seems to be decreased synteny towards the center of the island.

**Figure 2.**
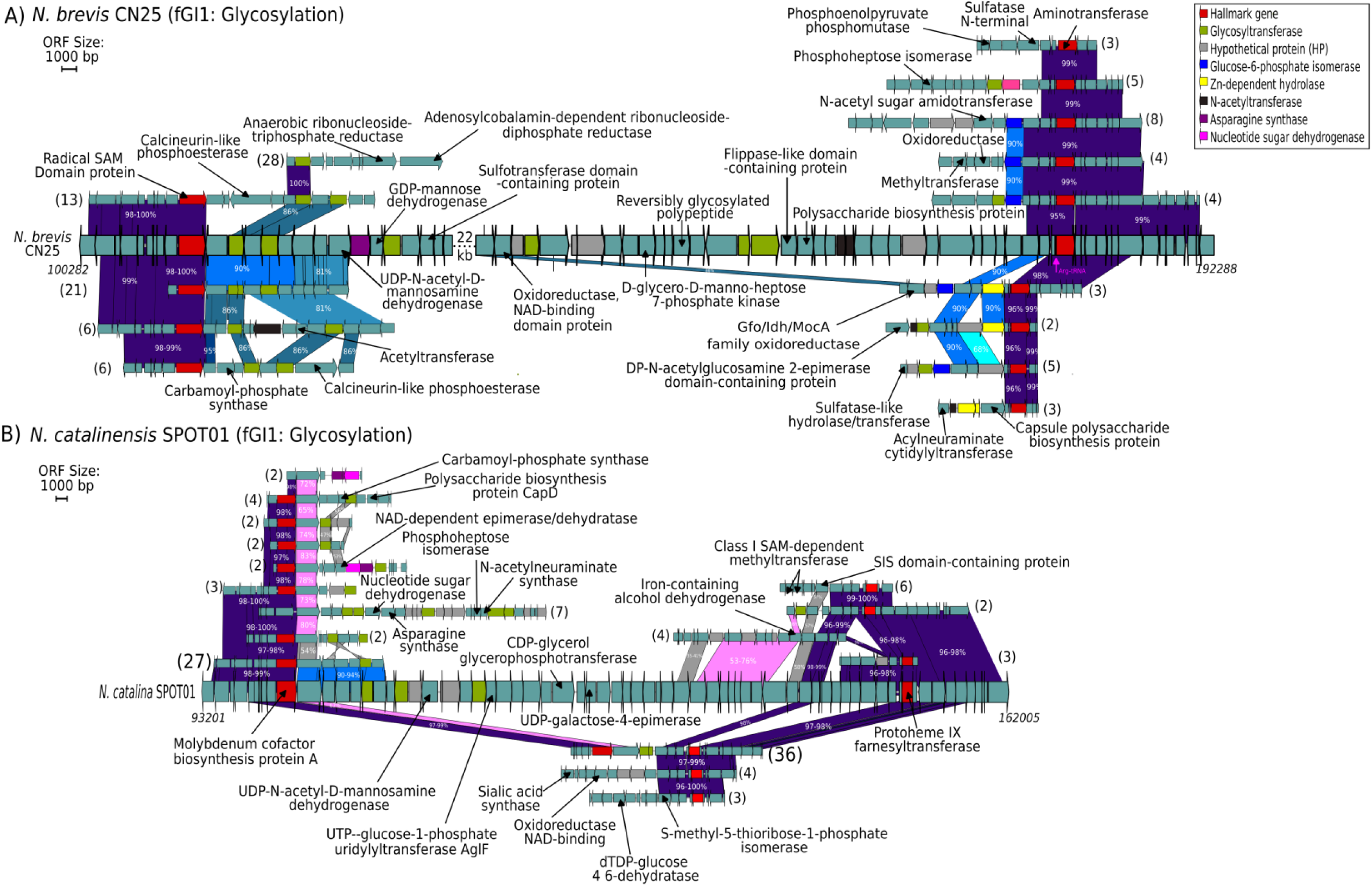
Flexible genomic islands (fGIs) related to glycosylation detected in *N. brevis* CN25 (A) and *N. catalinensis* SPOT01 (B). fGI versions and functional annotations of the ORFs are shown. The number in parenthesis indicates the LRs number associated with each fGI version size increased for the most abundant. For the genome of *N. brevis* CN25, the synteny is broken for improved viewability of the fGI (22 kb). Blue Arrow and letter show the location of tRNAs.

### Phosphorus-related genomic islands

Phosphorus is often the limiting nutrient in the open ocean photic zone and it has been shown to display major diversity in the gene clusters involved in phosphate and phosphonate transport (Molina-Pardines et al. 2023). It seems reasonable then, that a lot of evolutionary creativity is focused on the acquisition of such important nutrients. For *N. brevis* population, two genomic islands were related to the transport of phosphorus, with the addition of phosphonate (*phn* gene cluster, “fGI2”) and phosphate (*pst* gene cluster, “fGI3”) transporters (Figure 3A and 3B), and for *N. catalinensis* only one island, with the addition of a *pst* gene cluster (fGI3) (Figure 3C). In both cases, the GI versions found in the Pacific Ocean isolates were retrieved at high similarity (even if in smaller numbers), the alternative versions, larger and more complex, were found more frequently in the Mediterranean (with 51 and 70 LRs for phosphonate and phosphate transporters in *N. brevis* CN25 and 81 for phosphate transporter in *N. catalinensis* SPOT01). Given the scarcity of phosphorus in the Mediterranean and North Atlantic, it is to be expected that here lineages with more transporters (and likely higher affinity) to this nutrient would be more prevalent. However, since the Pacific Ocean versions were also retrieved in our Mediterranean population, some of these gene clusters seem to have a global distribution.

**Figure 3.**
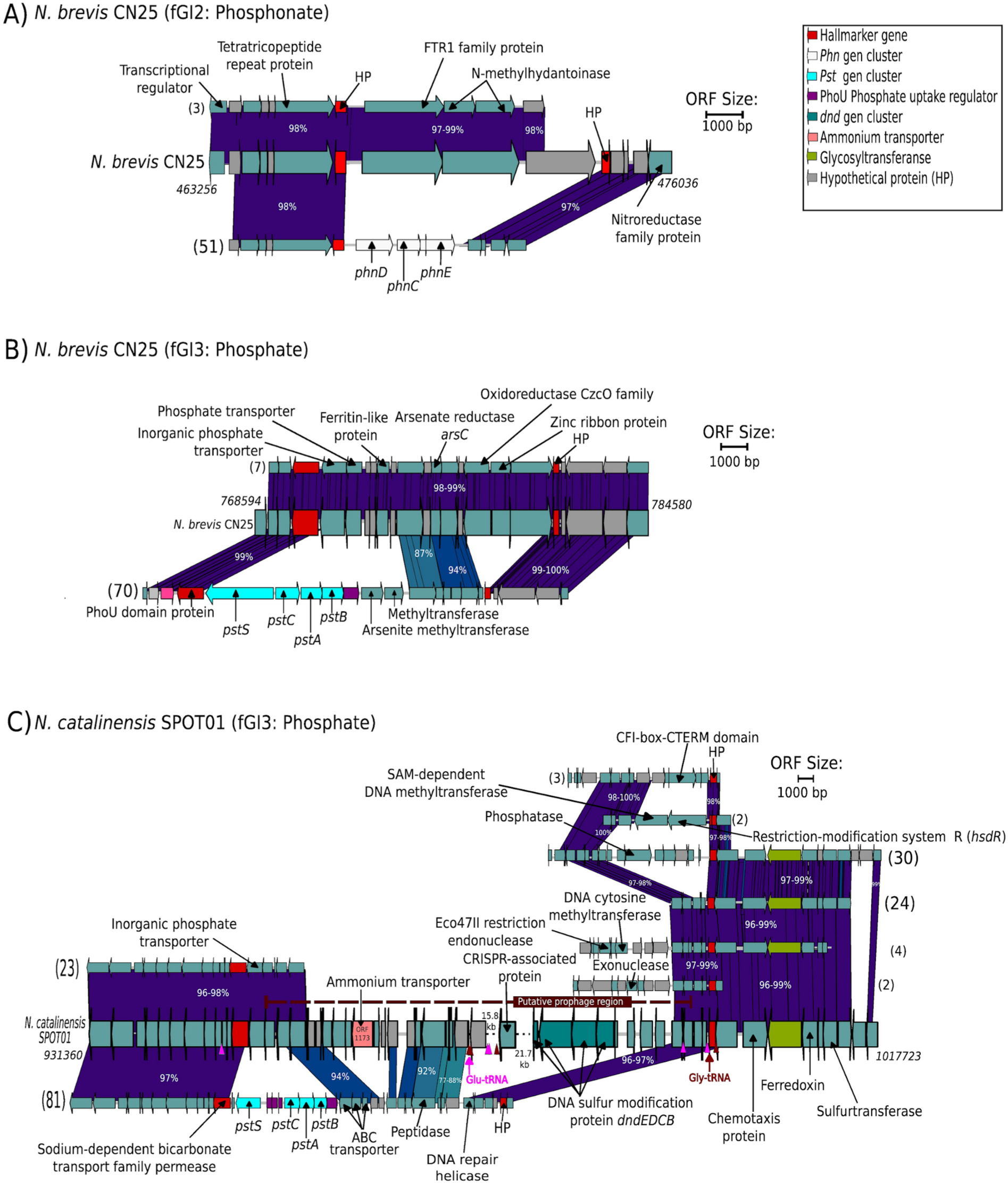
Flexible genomic islands (fGIs) related to phosphonate and phosphate in *N. brevis* CN25 (A, B), and to phosphate in *N. catalinensis* SPOT01 (C). fGI versions and functional annotations of the ORFs are shown. The number in parenthesis indicates the LRs number associated with each fGI version. For the genome of *N. catalinensis* SPOT01 the synteny is broken for improved viewability of the fGI (15.8 and 21.7 Kb) and the dotted line shows the predicted viral region by Ahlgren et al. 2017. Blue and maroon arrows indicate the location of Glu-tRNA and Gly-tRNA in the SPOT01 genome, while Blue and maroon triangles represent the alignment locations (average 14 nt) of Glu-tRNA and Gly-tRNA sequences within the fGI.

For SPOT01, a putative provirus has been suggested to be found next to the phosphorus island (Ahlgren et al. 2017). Some of the ORFs (NMSP_1215, NMSP_1217, NMSP_1226, NMSP_1228) predicted within this putative prophage in Ahlgren et al. 2017 and those shown to have hits to known viral proteins deposited databases were found among our Mediterranean metagenomic LRs (68 LRs found in the protein similarity analysis, Table A6). However, putative viral genes were not predicted for these LRs (Table A6) and a re- analysis of the strain SPOT01 genome using VirSorter2 (https://github.com/jiarong/VirSorter2) only detected an ORF associated with a viral gene (ORF_995) located far from the putative prophage region. Therefore, it appears that the prophage region was not present in our metagenome and/or that the SPOT01 viral region was misidentified. It is also interesting that in the Mediterranean *N. catalinensis* population, most long reads (81) have lost one (ORF_1173) of the two ammonium transporters present in the SPOT01 genome (see below and Figure 3C and Table A4). Additionally, a *dnd* gene cluster (From ORF_1233 to ORF_1236) involved in phosphorothionation, potentially involved in defense against phages was detected in the same region of the SPOT01 genome (Figure 3C and Table A4) that was absent in the Mediterranean population.

### Ammonium genomic islands

Some functions found in the flexible genomic islands as ammonium transporters were deleted in the *N. brevis* (fGI4) and *N. catalinensis* (see fGI3) populations (Figure 4 and Figure 3C). For *N. brevis*, the larger number of LRs (n=39) had an island version without the ammonium transporter (ORF 1313) present in the strain genome, while other versions (2 LRs) conserved this transporter or was replaced by a glycosyltransferase and an acetyltransferase (5 LRs) (Figure 4). For the *N. brevis* population in our metagenome, only one of the ammonium transporter annotated genes (ORF_1339, 1580 bp) was detected by recruitment from the core genome, (Table A3). Thus, it is likely that only one ammonium transporter is present in the Mediterranean population compared to the two identified in the Pacific genome. Similar results were found in *N. catalinensis* population, in which only one of the two ORFs (ORF_1173 and ORF_1559) detected as ammonium transporters in the SPOT01 genome recruited in our SR metagenomes (ORF_1559, 1568 bp, Table A4). Actually, in the *N. catalinensis* population, ORF_1173 was deleted in the LRs of the most common version of the phosphonate genomic island (Figure 3C and see above). Overall, ammonium transporters seem to be less prevalent and diverse in the Mediterranean than in the Pacific, which again fits with a limitation by phosphorus in the first water body.

**Figure 4.**
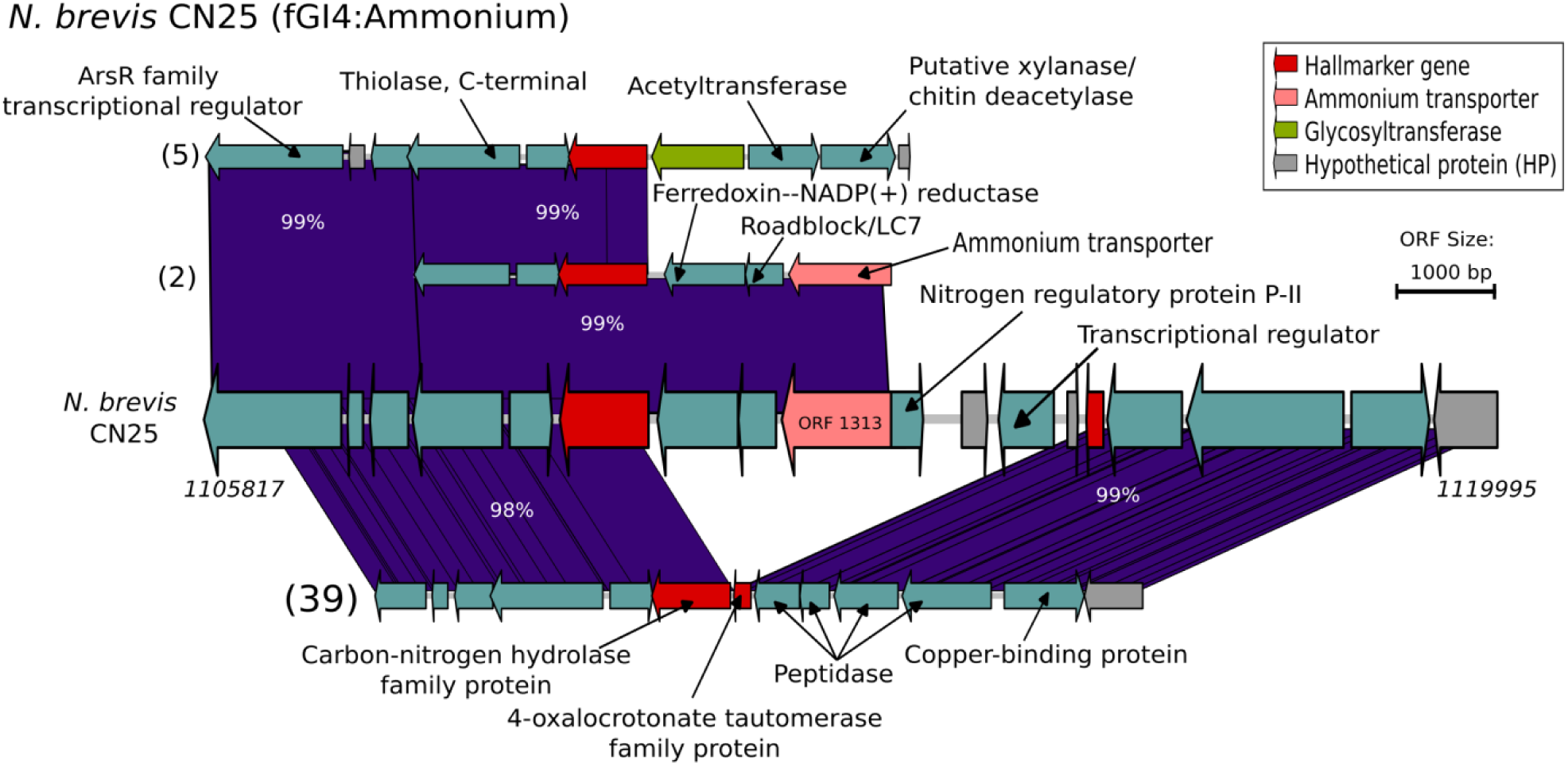
Flexible genomic islands related to ammonium in *N. brevis* CN25. fGI versions and functional annotations of the ORFs are shown. The number in parenthesis indicates the LR number associated with each fGI version. The most abundant GI version in CN25 showed the deletion of an Ammonium transporter.

**Figure 5.**
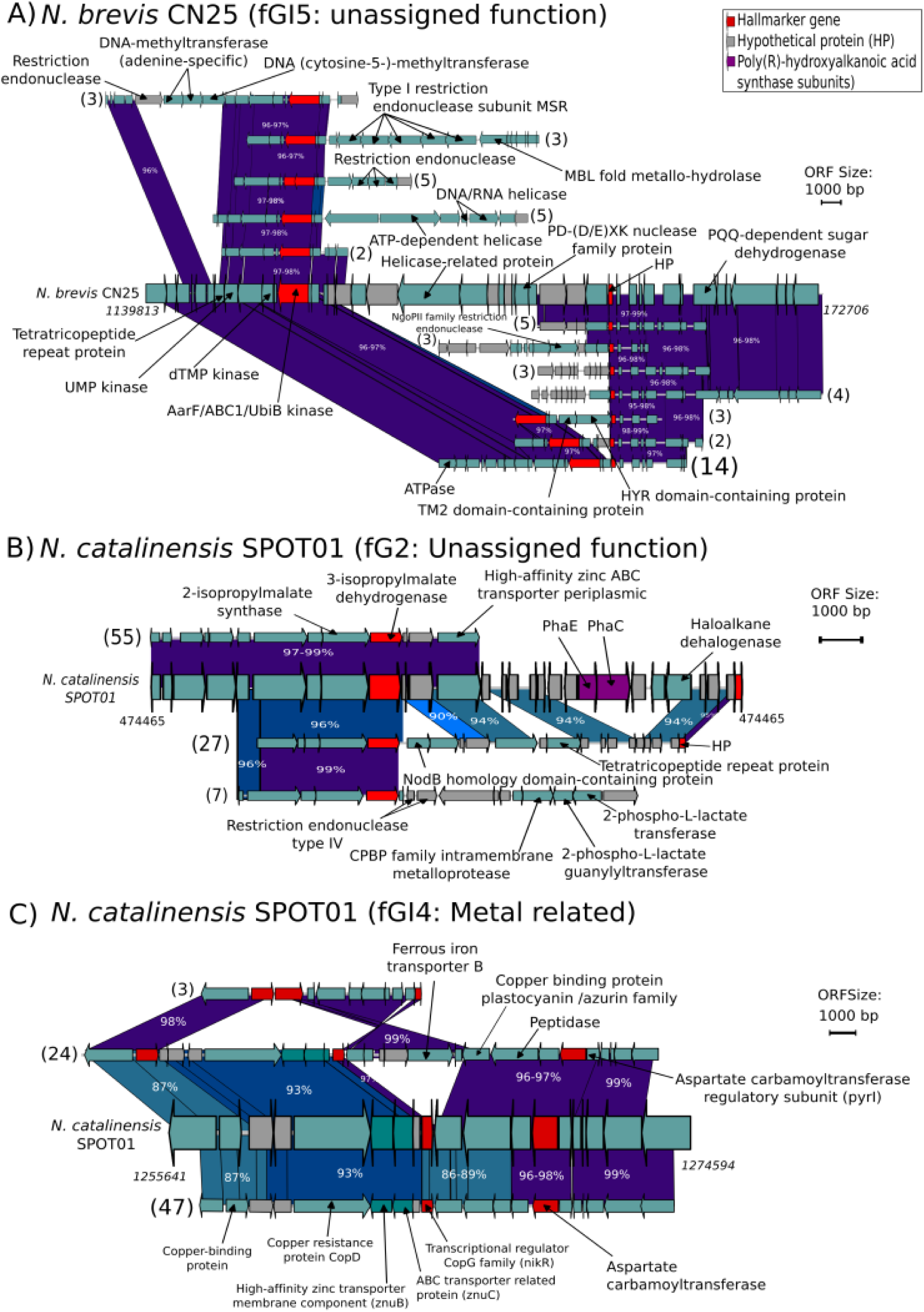
Genomic islands with unassigned function in *N. brevis* CN25 (A) and *N. catalinensis* SPOT01 (B), and a fGi related to metals. fGI versions and functional annotations of the ORFs are shown. The number in parenthesis represents the LR number associated with each fGI version. For the fGI in *N. brevis* CN25 (A), a possible defensive function is suggested.

### Metal related and unassigned fGIs

One metagenomic island with genes with no defined function was found for each of the reference genomes CN25 and SPOT1 respectively (Figure 4). For *N. brevis* population, fGI4 (Figure 4A), had 12 versions delimited by the hallmark genes. The most frequent (>14 LRs) had a deletion of the CN25 island (the hallmark genes appeared contiguous) while other versions had several restriction endonucleases and DNA-related proteins like helicases (Figure 4A) together with some methyltransferases which suggest that this fGI has a defense function. For the genome of *N. catalinensis* SPOT01, a single CRISPR- associated protein (ORF_1212) was detected but no tandem repeats or spacers (Figure 4B). However, the large number of hypothetical proteins prevents assigning reliably any biological role to the gene clusters as already reported (Santoro et al, 2015).

The metagenomic island labelled as fGI2 in the SPOT1 genome was also very hard to interpret at the level of biological role. Most LRs in the Mediterranean population were associated with a version similar to the Pacific population (at least for the three ORFs after the hallmark gene). Another version (27 LRs) has a completely different gene complement with the loss of the two poly(R)-hydroxyalkanoic acid synthase subunits *phaC* and *phaE*, found in SPOT1 and addition of genes related with NodB, Formyl transferase, and tetratricopeptide repeat protein (Figure 4B). Another GI version (7 LRs) had genes associated with restriction endonuclease, stress response translation, and CPBP family intramembrane metalloprotease.

For SPOT1, another underrecruiting region (fGI4) was found with a copper binding protein located nest to genes associated with copper (resistance or other binding proteins, *cop* operon) (Figure 4C). A high affinity for copper coupled to potential toxicity from this metal has been reported for isolates of *Nitrosopumilus maritimus* strain SCM1 (Shafiee et al., 2021). This could be the reason why in *N.* catalinensis population we found an fGI with numerous genes encoding copper-binding proteins and associated copper resistance genes. In the Mediterranean population of *N. catalinensis*, most of LRs covering the region (n=24) had an insertion of a ferrous iron transporter among other genes within the copper- related genes (Figure 4C). Still, the most common version (n=47) was syntenic with the Pacific isolate albeit at lower (ca. 85%) similarities. the island version with genes of copper-binding proteins and copper resistance and with deletion of ferrous iron transporters, which was present in another island version with half of LRs (n=24). The last island version (n=3) had the deletion of the copper resistance genes and the ferrous iron transporter inverted in relation to the most common version mentioned above (Figure 4C). The growth of the Isolated *N. maritimus* strain SCM1 was determined by Fe availability, and the lack of siderophore production and the use of exogenous siderophores for Fe uptake have been reported (Shafiee et al., 2019). We hypothesized that the presence of a ferrous iron transporter in some LRs (n = 24) of the *N. catalinensis* population could be associated with iron uptake from the environment, while for LRs without this iron transporter (n = 47), the use of exogenous siderophores like *N. maritimus.* with the iron acquisition strategy in the Mediterranean population of *N. catalinensis*. Siderophore production was not found in the genomes of SPOT1 or CN25 (analysis by AntiSMASH v7, data not shown).

## DISCUSSION

Our objective here has been to identify flexible genes in populations (cells of the same species present in a single place and time) of chemolithotrophic marine archaea. Since the retrieval of several cultures from these hard-to-grow microbes does not seem feasible, we have used metagenomic recruitment from Illumina (SR) metagenomes to delimit high variability regions that presumably are gene clusters variable from one cell to another and thus underrecruiting Illumina short reads. With this approach, it is only possible to detect genomic regions (islands) subjected to this kind of sequence diversity or flexible genomic islands. A similar approach was used before for detecting variable regions in populations of extremophilic archaea (Cuadros-Orellana et al., 2007) and bacteria (Pašić et al. 2009). Revealing high diversity in their populations. Recently, studies of read diversity in metagenomes and pure cultures indicated also a high intrapopulation diversity for these extremophilic heterotrophs (Viver et al. 2024). AOA on the other hand are marine chemolithoautotrophic microbes with simple requirements in terms of nutrients. Our results indicate that they indeed have much less diversity. Judging by the glycosylation island, which tends to be lineage-specific, in our samples there were likely less than a dozen different coexisting clonal lineages. Still, we detected diversity in regions of likely ecological importance such as those related to the transport of the two major limiting nutrients: phosphorus and ammonia. The numbers and diversity of these transporters were consistent with the measured values and limiting role played by phosphorus in the two oceanic regions (Pacific and Mediterranean). If the latter is so limited by this nutrient, it seems reasonable that fewer requirements of high-affinity ammonia transported are needed since growth would be limited by phosphorus anyway.

In a recent study, some of us have studied the diversity of O-chain polysaccharide genomic islands in Pelagibacterales (heterotrophic bacteria) and have found a very large diversity, reaching easily 130 different gene clusters in a single population (single species single sample) (Haro-Moreno et al, 2024). This is in sharp contrast to the results described here for a chemolithotrophic archaeon. Although it is still early to conclude, the results fit with the diversity of major nutrients in both cases (dissolved organic matter in one versus ammonia and phosphorus in the other). Thus, the difference in genomic diversity within populations reflects as expected the diversity of the major substrates for each microbe.

The approach we have used here opens a new path to explore the flexible genome of uncultured microbes by the use of long-read metagenomes that can be leveraged to assemble significant contigs of these regions that very seldom (if at all appear in short read metagenome-assembled genomes (MAGs). SR metagenomes assemble very poorly at regions that are variable in the population. Furthermore, the standard methods used to assess the completion of MAGs use core genes that cannot be used to estimate how much of the flexible genome is recovered. Although we have recovered only small fragments due to the condition of the presence in the metagenomic read of the hallmark gene, it is possible to retrieve much larger fragments by overlapping long reads at high similarity and overlap (e.g. 1,000 nucleotides overlap). This would allow the reconstruction of important gene clusters such as those implicated in the biosynthesis of bioactive compounds or pathogenicity islands. This would enlarge the already gigantic scale of the known sequence space of prokaryotes.

## EXPERIMENTAL PROCEDURES

### Sampling, Quality Filters and Sequence Analysis

A water sample was collected on October 8, 2021, to a depth of ∼75 m in the western Mediterranean Sea (37.35361◦N, - 0.286194◦W). Previous work determined the lower photic (LP) zone in this depth with a high abundance of AOA (Haro-Moreno et al. 2018). The LP sample was subsequently filtered for different pore filter polycarbonate filters (20-, 5-, and 0.22-µm Millipore) [see details in Haro-Moreno et al. (2021)] and gDNA was extracted using the 0.22-µm filter and the genomic tips Kit (Qiagen) following the manufacturer’s instructions. Metagenomes were sequenced using Illumina Nextseq (150 bp, paired-end reads) (Novogene, United Kingdom) and PacBio Sequel II (one 8M SMRT Cell Run, 30-h movie) (Novogene, United Kingdom). The quality of the Illumina reads was checked with FastQC version v0.11.9 (Andrews 2010) and low-quality bases (per base sequence quality <33) were removed with Trimmomatic version 0.39 (Bolger et al. 2014). Highly Accurate Single-Molecule Consensus Reads (CCS reads) were generated using the CCS v6 program of the SMRT- link package (PacBio, Menlo Park, CA, USA) with a minimum number of full-length subreads of 15 (99.95 base call accuracy).

### Microbial community structure and AOA phylogenetic diversity from the LP zone

To assess the relative abundance of the phylum Thaumarchaeota in the lower photic zone, 16S rRNA gene reads were retrieved from Illumina (Short Read “SR”) and PacBio (Long Read “LR”) sequences. A subset of 20 million reads of the SR library was extracted using vsearch v2.14.1 (Rognes et al. 2016) and after were aligned against a nonredundant version of the SILVA database v138 with an E-value <10−5. Sequences that matched this database were aligned against archaeal and bacterial 16S rRNA hidden Markov models (HMMs) using ssu-align 0.1.1 (github.com/EddyRivasLab/ssu-align). For the LRs, 16S rRNA sequences were extracted using barrnap v0.9 (github.com/tseemann/barrnap) from total PacBio CCS15 reads. The 16S rRNA sequences obtained were classified using the sina algorithm (Pruesse et al. 2012) against the SILVA taxonomy database.

To analyze the phylogenetic diversity of the phylum Thaumarcheota on the low photic zone, the 16S rRNA gene extracted from the total PacBio CCS reads and that belonging to Thaumarchaeota were clustered with 97% similarity cut-offs using CD-HIT (Fu et al. 2012), and along with 16S rRNA gene sequences extracted of reference genomes from GenBank and IMG databases (Table A7) by barrnap v0.9 (github.com/tseemann/barrnap), were aligned using muscle v3.8.1551 (Edgar 2004). *Sullfolobus solfataricus* SULA and *Thermoproteus tenax* Kra 1 sequences were used as outgroups. The alignment was submitted to JModelTest v.2.1.10 (Darriba et al. 2012) which determined that the GTR + G +I model was the most appropriate substitution model. The phylogenetic estimation was performed using trimAl v1.4.rev15 (Capella-Gutiérrez et al. 2009) with default parameters and bootstrapping of 10,000. The tree was visualized with iTOL v6.81 (Letunic and Bork 2019).

### Metagenomic read recruitments

Our strategy to obtain data on the local pangenome of the abundant species of Thaumarchaea was based on a differential recruitment approach. Briefly, using a reference genome that is assumed to be present in the sample, the regions that recruit significantly fewer reads are considered as “metagenomic islands’’ i.e regions of the reference genome with little or no presence in the metagenome and likely belonging to the flexible genome of the species (Rodriguez-Valera et al. 2016). As a first step, we identified the marine microbes populations belonging to phylum Thaumarcheota that were present in the lower photic zone. Genomes of phylum Thaumarcheota reported in the literature were used to recruit reads from our metagenomics Illumina read datasets from the LP sample. The normalized RPKG value (number of reads recruited per kb of genome per Gb of metagenome) and coverage values (alignment of Illumina metagenomic reads in a subset of 40 million reads) were calculated for all genomes using BLASTN (Altschul et al. 1997) using a cutoff range from 95 to 99% nucleotide identity over a minimum alignment length of 50 nucleotides. Genomes with the highest RPKG values were selected to discover genomic islands and construct recruitment plots using the R package ggplot2 (Ginestet 2011). Metagenome-assembled genomes (MAGs) belonging to phylum Thaumarchaeota were not included in the recruitment approach and GIs searching due to low recovery of the flexible genome (Rodriguez-Valera et al. 2009; Haro-Moreno et al. 2021).

### Identification and analysis of flexible genomic islands (fGIs) in Thaumarchaeota populations

The genomic islands were detected using two approaches: a) The reference genomes with the highest RPKG values were annotated using Prodigal v2.6.3 (Hyatt et al. 2010) and the open reading frames (ORFs) obtained from this analysis, were recruited against the Illumina metagenomic dataset from LP zone, and metagenomes published in our previous study in water samples from the same Mediterranean sea site (Haro-Moreno et al., 2018). Those zones in the ORF recruitment that did not recruit evenly ≥ 10% of the mean value of RPKG by metagenomic dataset, and b) those gaps or under-recruiting regions detected in the recruitment plots, were considered as candidate genomic islands. For the identification of the genetic variation in candidate GIs, the ORF of 1 upstream and 1 downstream of each GI in the reference genome (called hallmark gene), and those ORFs located inside of GI (called ORF-inside) were extracted and aligned against the PacBio CCS reads from the LP sample (See Figure A2-A6 for details). The LRs that had >95% of sequence identity with a hallmark gene and those with > 50% with ORF-inside were chosen for the search of GI versions in their respective AOA populations. The LRs shared between the hallmark gene and ORF-inside were obtained using custom command-line tools. The LRs unique to the hallmark gene with positive strand to the inside of the island were analyzed for the detection of new variation (called ORF-local, ORFs that are not in the reference genome, however, they are in the local microbial population). The LRs were annotated using Prodigal (Hyatt et al. 2010), and the synteny of their ORFs and those obtained from the reference genomes were visualized using the flex tool (github.com/asierzaragoza/flex2). Functions of predicted ORFs were manually curated and revised by a comparison of sequence homology against the NCBI NR database using DIAMOND (Buchfink et al. 2014) and the UniProt databases using BLASTp (Altschul et al. 1997).

## AUTHOR CONTRIBUTIONS

**Francisco Rodriguez-Valera**: Conceptualization, funding acquisition; writing–original draft. **Pablo Suárez-Moo:** formal analysis; data curation; writing–original draft. **Jose M. Haro-Moreno:** formal analysis; data curation.

## ACKNOWLEDGEMENTS

This work was supported by grant “FLEX3GEN” PID2020-118052GB-I00 (cofounded with FEDER funds) from the Spanish Ministerio de Economía, Industria y Competitividad to Francisco Rodriguez Valera and Jose Manuel Haro Moreno. was supported with a PhD fellowship from Margarita Salas program, cofounded by the Spanish Ministerio de Universidades and the European Union—Next-Generation EU (2021/PER/00020).

## DATA AVAILABILITY STATEMENT

Metagenomic datasets have been submitted to NCBI SRA and are available under BioProject accession number PRJNA1088973.

