## Supplementary figures for "Microdiversity in marine pelagic ammonia-oxidizing archaeal populations"

**SUPPORTING INFORMATION**

Figure A1. Microbial community structure and phylogenetic diversity of the phylum Thaumarcheota in the metagenomes of the lower photic zone. A) Relative abundance of phyla based on 16S rRNA gene fragments of Illumina (SR) and PacBio CCS15 (LRs) metagenomic reads. The phylum Proteobacteria was divided into class-level classifications. Only those groups with abundance values larger than 1% are shown. B) Phylogenetic tree based on 16S rRNA gene LRs, bold numbers in parentheses represent the number of sequences by cluster group obtained in the dereplicate process (97% similarity). The NCBI accession of the reference sequences is shown in parentheses. The color red shows the AOA species used in the genomic island analysis.

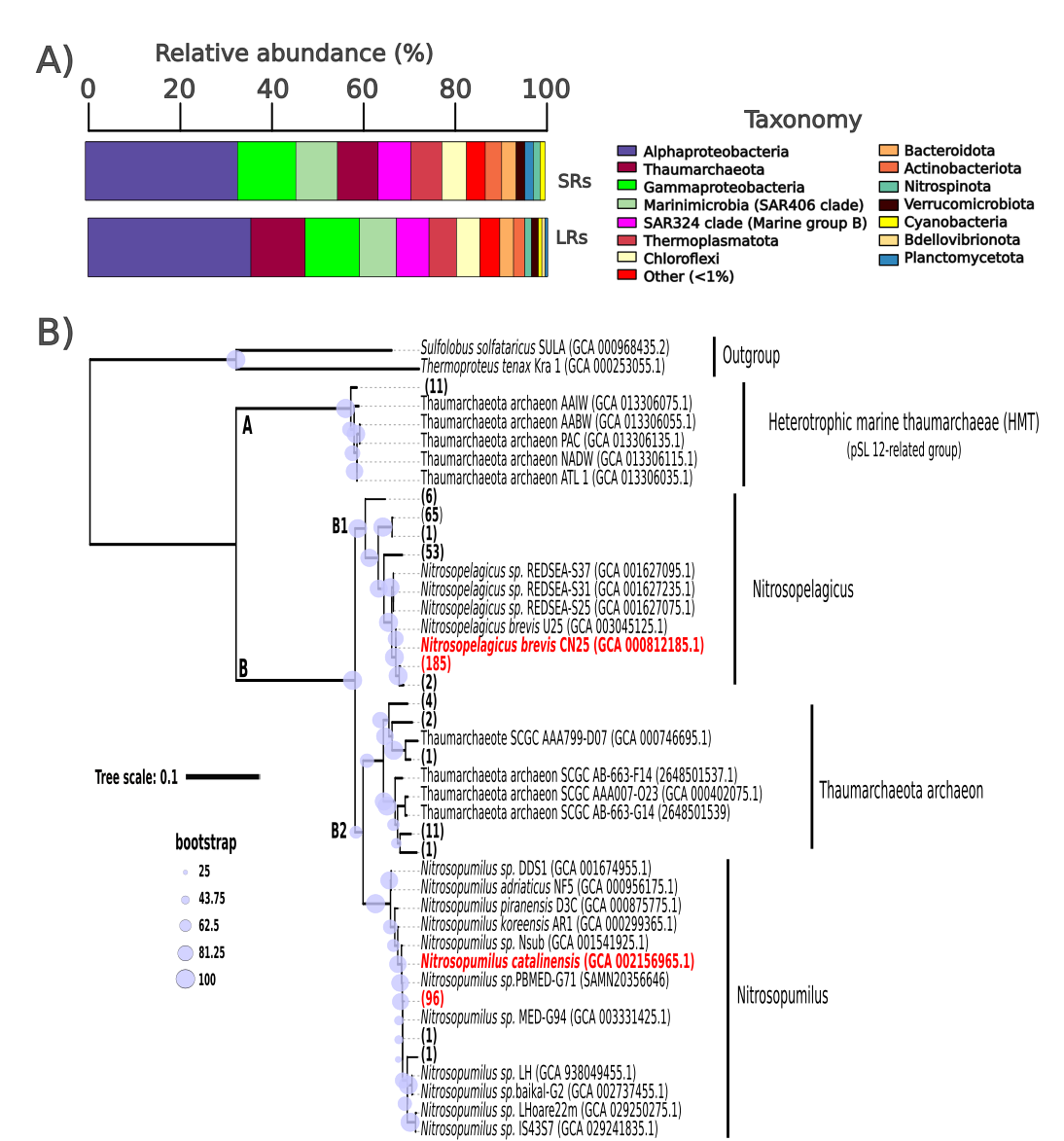

Figure A2. Exploration of the genetic variation within the GIs. LR CCS15 reads corresponding to a specific GI version were overlapped to generate the longest LR CCS15 read (with 97-99% sequence identity and 1,000 bp alignment). The hallmark genes (red color), CDS-inside (pink color), and CDS-local (green color) are shown. The LRs that had >95% of sequence identity with a hallmark gene and those with > 50% with ORF-inside were chosen for the search of GI versions in their respective AOA population. ORF-local are ORFs that aren't in the reference genome, however, they are in the local microbial population and were blasted against the LR metagenome (>95% sequence identity) to discover other GI versions and numbers of LR associated with them.

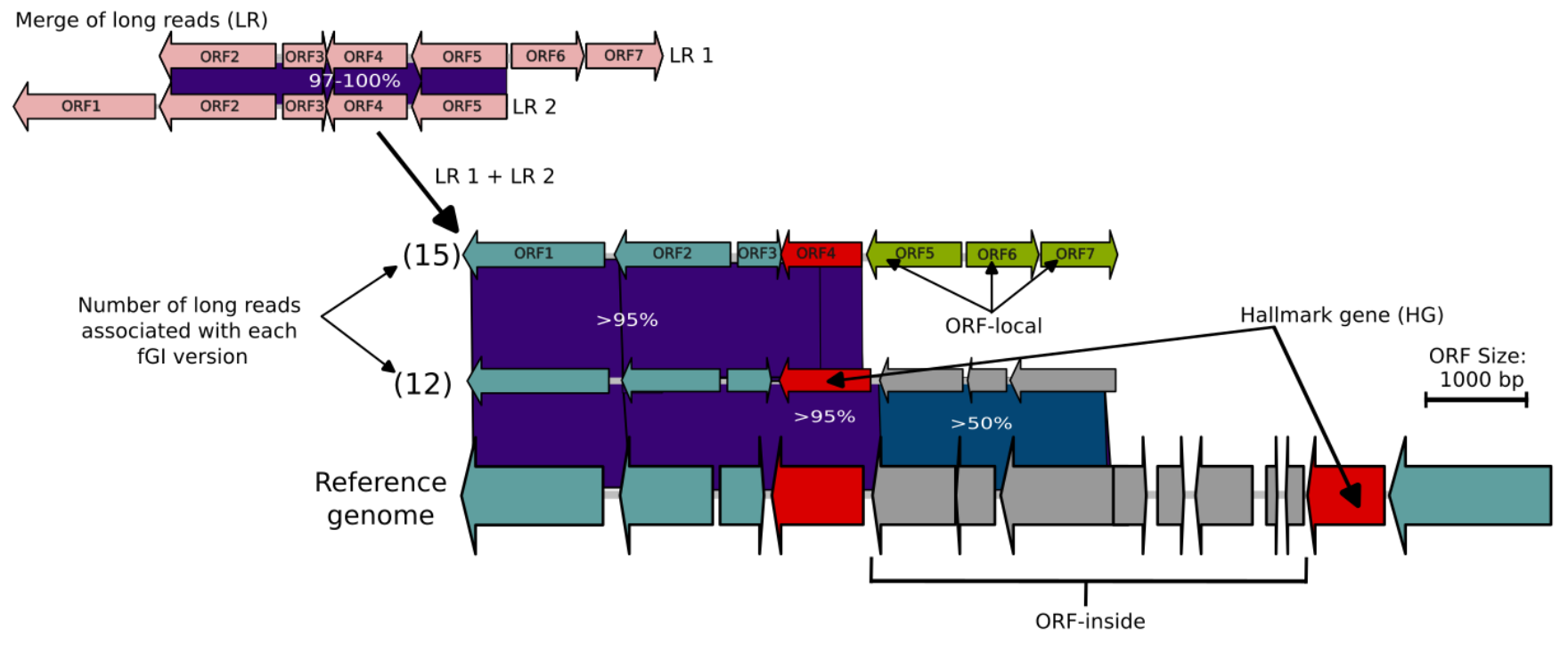

Figure A3. fGI versions related to glycosylation (A), phosphonate (B) and phosphate (C) clusters in *N. brevis* CN25. The number in parentheses indicates the total number of LRs and those used for the merge of long reads for each fGI version. The vertical dashed line separates the right and left sides of fGI1.

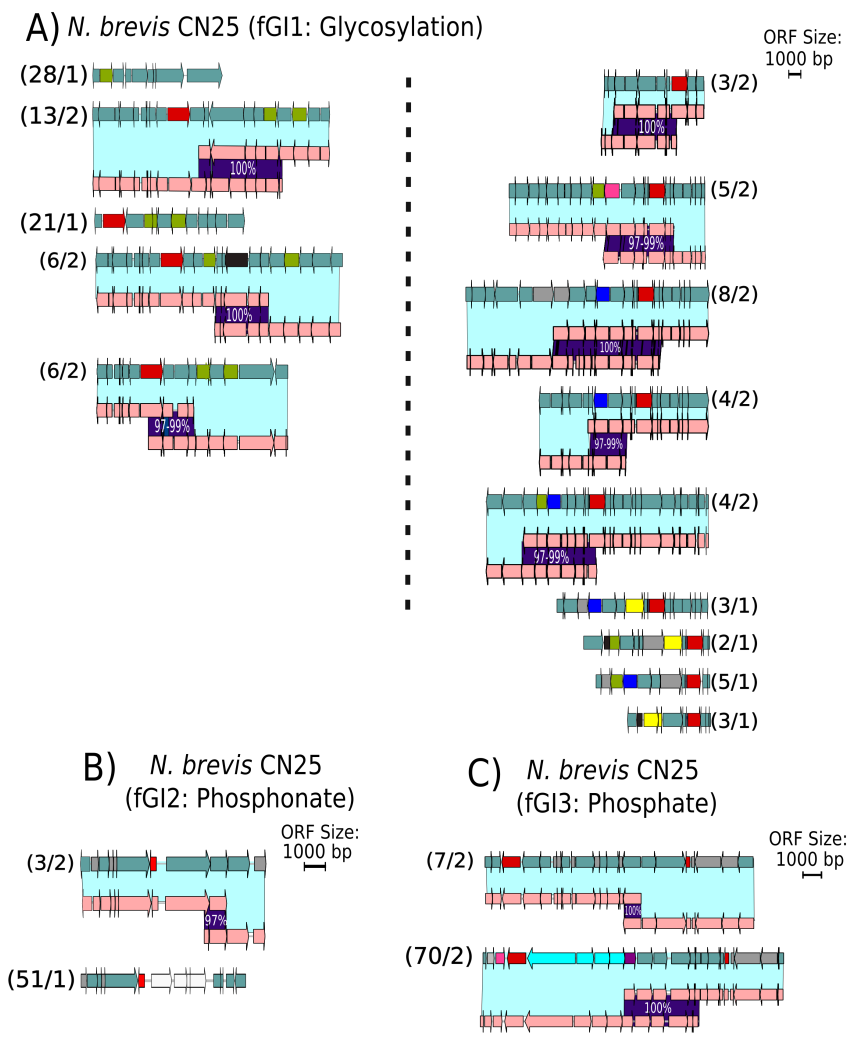

Figure A4. fGI versions related to ammonium (A) and unassigned function (B) in *N. brevis* CN25. The number in parentheses indicates the total number of LRs and those used for the merge of long reads for each fGI version. The vertical dashed line separates the right and left sides of fGI5.

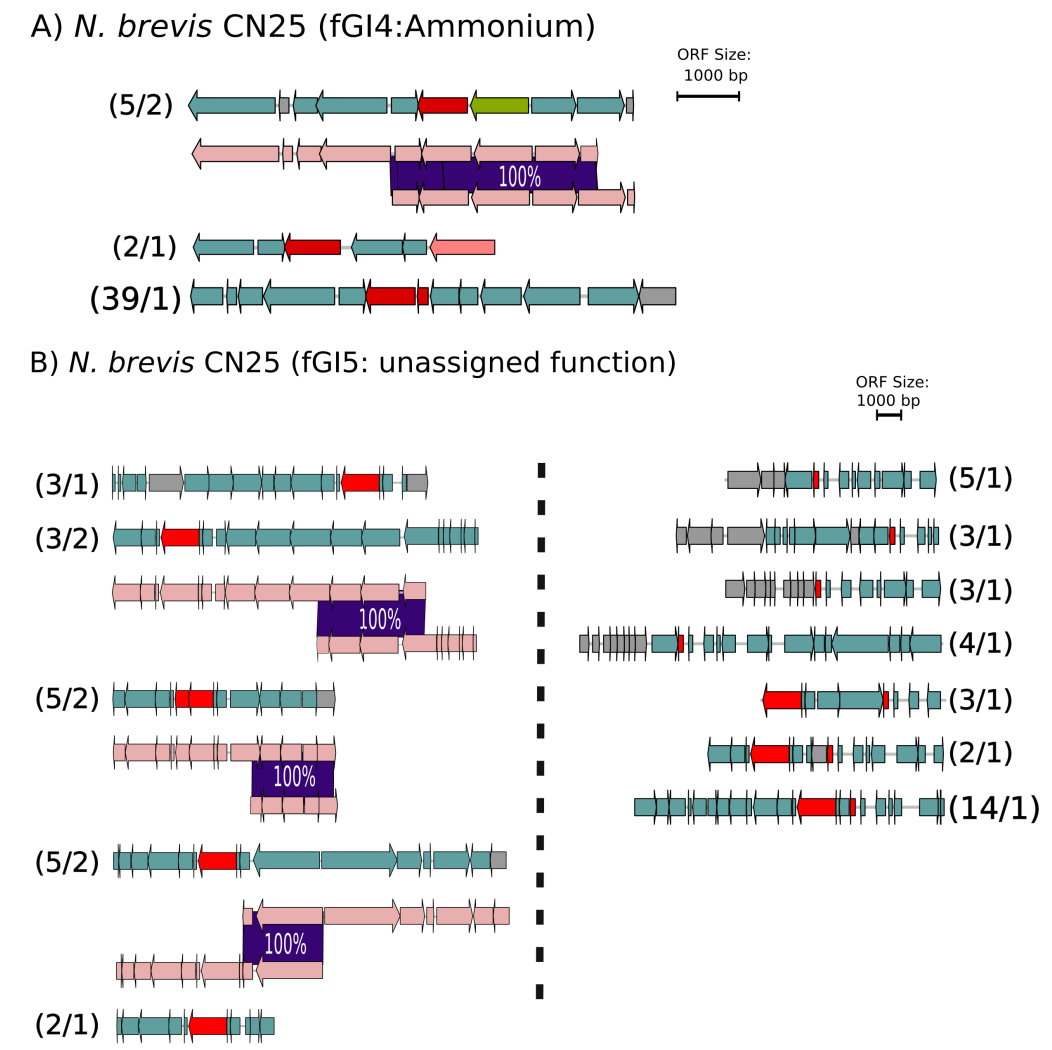

Figure A5. fGI versions related to glycosylation in *N. catalinensis* SPOT01. The number in parentheses indicates the total number of LRs and those used for the merge of long reads for each fGI version. The vertical dashed line separates the right and left sides of fGI1.

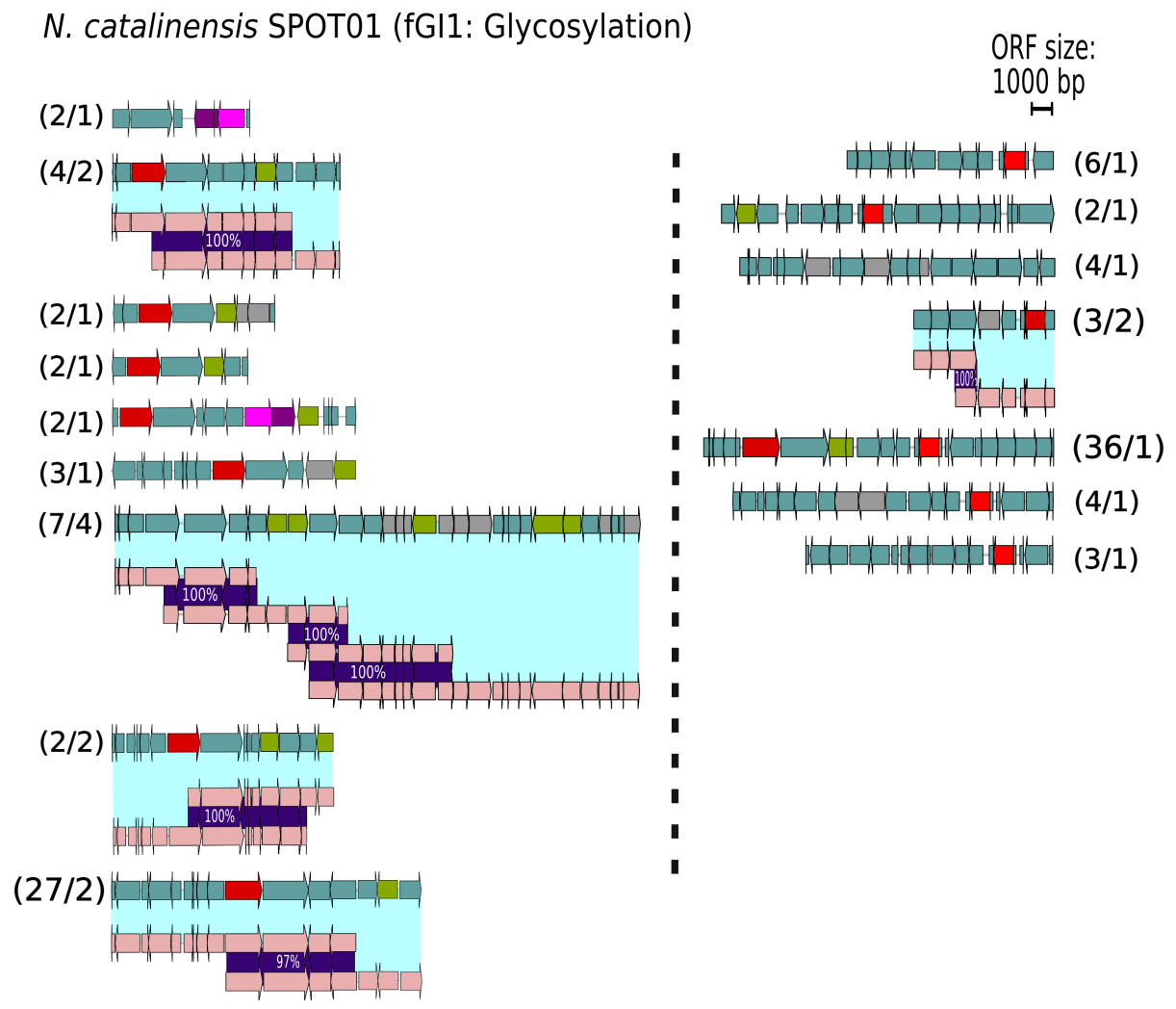

Figure A6. fGI versions related to phosphate (A), Unassigned function (B), metal related (C) in *N. catalinensis* SPOT01. The number in parentheses indicates the total number of LRs and those used for the merge of long reads for each fGI version. The vertical dashed line separates the right and left sides of fGI5.

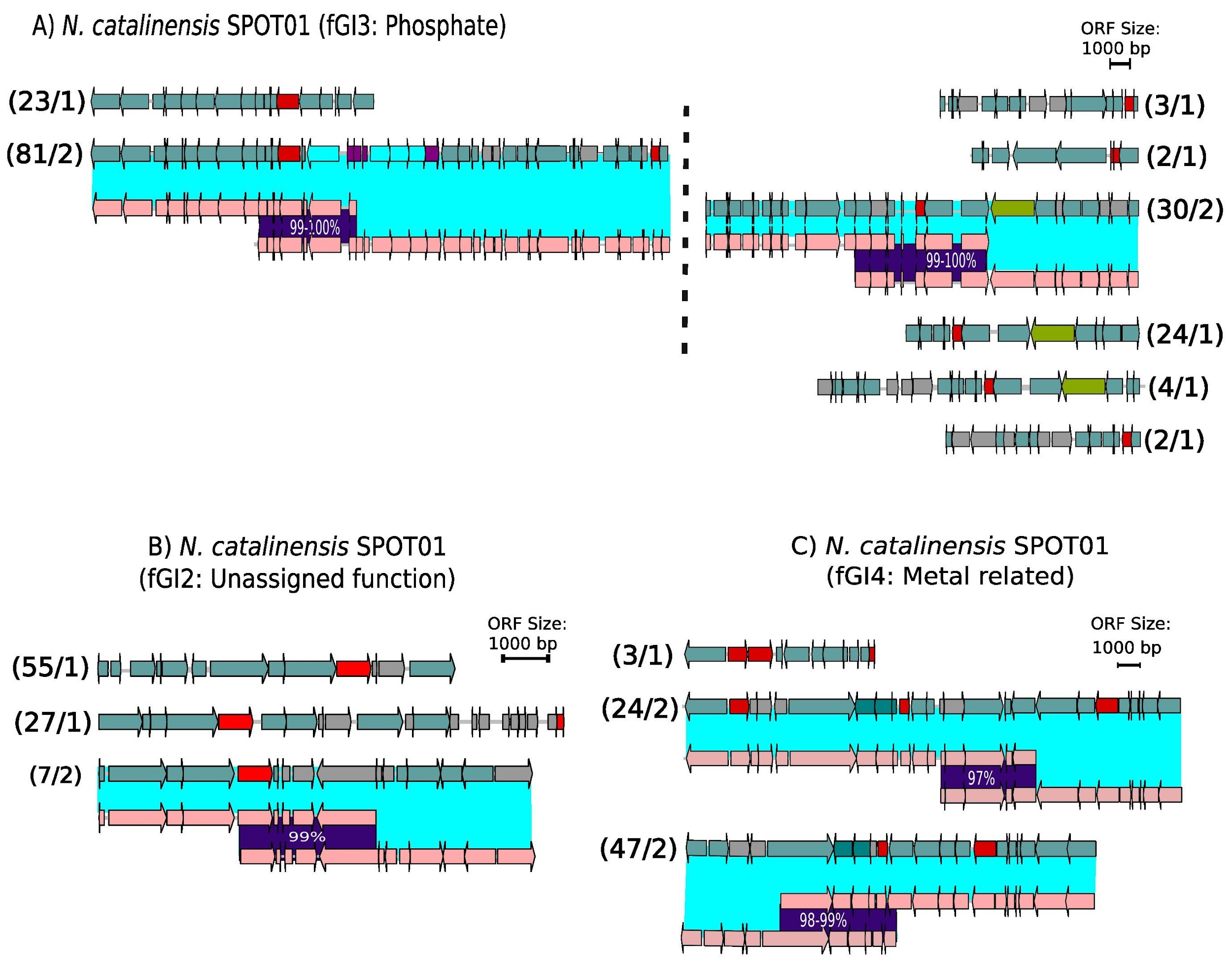

Table A1. Summary statistics of the short-read and long-read sequencing obtained from lower photic zone.

Table A2. Reference genome recruitment against the short-reads from the lower photic zone. RPKG and coverage values are shown in a range of identity percentages from 95 to 99%. Classification by GTDB-tk and the sample and sequencing characterization are also shown. The color red shows the AOA species used in the genomic island analysis.

Table A3. Annotation and recruitment of the Nitrosopelagicus brevis CN25 genome. The functional annotation of ORFs by comparison against NCBI NR and Uniprot databases is shown. The genome CN25 was recruited against the SR metagenome and previous metagenomic datasets from the off-shore Western Mediterranean. ORFs associated with the GIs (yellow), hallmark gene (red), S-layer (Blue), Ammonia monooxygenase (AMO) (bright green), and ammonium transporter (Turquoise) are shown. The tRNAs are highlighted in green.

Table A4. Annotation and recruitment of the Nitrosopumilus catalinensis SPOT01 genome. The annotation of ORFs by comparison against NCBI NR and Uniprot databases are shown. The genome SPOT01 was recruited against the SR metagenome and previous metagenomic datasets of off-shore Western Mediterranean. ORFs associated with the GI (yellow), hallmark gene (red), S-layer (Blue), Ammonia monooxygenase (AMO) (bright green), and ammonium transporter (Turquoise) are shown. The tRNA are shown in green

Table A5. Annotation of the Nitrosopumilus maritimus SCM1 genome. The annotation of ORFs by comparison against NCBI NR and Uniprot databases are shown. ORFs associated with S-layer and tRNAs are shown in blue and green, respectively.

Table A6. LRs associated with the putative provirus region suggested to the phosphorus island in N. catalinensis SPOT01 in Ahlgren et al. 2017. Protein similarity and results of the analysis of putative viral genes using VirSorter2 are shown.

Table A7. Reference genomes used in the phylogenetic analysis of the AOA in the Lower photic zone. NCBI Accession, strain, habitat, and source are shown. The color red shows the AOA species used in the genomic island analysis.
